# Patient induced pluripotent stem cell-derived hepatostellate organoids establish a basis for liver pathologies in telomeropathies

**DOI:** 10.1101/2021.11.19.469258

**Authors:** Young-Jun Choi, Melissa S. Kim, Joshua H. Rhoades, Nicolette M. Johnson, Corbett T. Berry, Sarah Root, Qijun Chen, Yuhua Tian, Rafael J. Fernandez, Zvi Cramer, Stephanie Adams-Tzivelekidis, Ning Li, F. Brad Johnson, Christopher J. Lengner

## Abstract

Patients with dyskeratosis congenita (DC) and related telomeropathies resulting from premature telomere dysfunction suffer from multi-organ failure. In the liver, DC patients present with nodular hyperplasia, steatosis, inflammation, and cirrhosis. We model DC liver pathologies using isogenic human induced pluripotent stem (iPS) cells harboring a causal DC mutation in *DKC1*, or a clustered regularly interspaced short palindromic repeats (CRISPR)-Cas9-corrected control allele. Differentiation of these iPS cells into hepatocytes or hepatic stellate cells followed by generation of genotype-admixed hepatostellate organoids revealed a dominant phenotype in the parenchyma, with DC hepatocytes eliciting a pathogenic hyperplastic response in stellate cells independent of stellate cell genotype. Pathogenic phenotypes could be rescued via suppression of AKT activity, a central regulator of MYC-driven hyperplasia downstream of *DKC1* mutation. Thus, isogenic iPS-derived admixed hepatostellate organoids offer insight into the liver pathologies in telomeropathies and provide a framework for evaluating emerging therapies.

## Introduction

Dyskeratosis congenita (DC) is a telomere biology disorder (telomeropathy) initially characterized by a clinical ‘triad’ of pathologies including nail dystrophy, oral leukoplakia, and abnormal skin pigmentation, now understood to be associated with bone marrow failure. DC frequently presents with additional pathologies including lung and liver fibrosis, intestinal barrier failure and inflammation, and osteopenia ^1–3^. DC is caused by mutations in various genes encoding proteins involved in telomere capping or elongation, most commonly the X-linked *DKC1* gene. *DKC1* encodes dyskerin - an integral component of the telomerase ribonucleoprotein complex important for maintaining sufficient telomerase activity ^4,5^. DC patients with loss-of-function mutations in *DKC1* have significantly shorter telomeres than age-matched controls ^6^, and isogenic human pluripotent cell-based models harboring causal DC mutations in *DKC1* exhibit attenuated telomerase enzymatic activity, shorter telomeres, and hallmarks of telomere dysfunction ^3,7,8^. Although dyskerin has additional roles in RNA pseudouridylation and ribosomal function ^2^, its dysfunction in telomere elongation is causal in DC, as the DC phenotypes caused by *DKC1* mutations are highly similar to those observed in DC cases caused by mutations in at least ten other genes whose shared function is telomere maintenance. Furthermore, any ribosomal defects are mild and are not correlated with the severity of DC phenotypes ^9^ and, moreover, restoration of telomere capping is sufficient to rescue DC phenotypes ^3,10–12^.

Unlike highly proliferating tissues such as the bone marrow and intestinal epithelium which exhibit stem cell failure in response to critical telomere shortening, lower turnover tissues such as the lung and liver present primarily with fibrosis ^4^. In the liver, DC patients frequently exhibit inflammation, steatosis, and nodular regenerative hyperplasia, all of which are often linked to cirrhosis ^13–15^. Several human studies have also underscored the frequency of telomerase mutations and critically short telomeres in otherwise idiopathic cirrhosis patients ^16,17^ Consistent with a causal role for telomere dysfunction in these liver phenotypes, telomerase RNA component (*Terc*) knockout mice normally exhibit cirrhosis upon carbon tetrachloride injury, and in this model adenoviral delivery of Terc is sufficient to reactivate telomerase, inhibit telomere shortening, and prevent cirrhosis initiation ^18^. Thus, cumulative evidence indicates that telomere failure in the liver is causal for the pathologies associated with DC, and potentially more broadly in idiopathic cirrhosis patients without a diagnosed telomeropathy.

Currently, there are no therapeutic options addressing the liver phenotypes associated with DC due to a lack of understanding of the cellular basis and molecular mechanisms underlying the phenotypes downstream of telomere dysfunction. Interestingly, in the fibrotic livers of cirrhosis patients, telomere shortening and senescence are limited to hepatocytes and are not observed in non-parenchymal cell types such as stellate cells and lymphocytes ^19^, suggesting that progression of cirrhosis may be caused by hepatocyte-specific telomere dysfunction.

Recent findings indicate that a population of TERT-high hepatocytes repopulate liver both in response to injury and during hepatic homeostasis ^20^. Experiments in human embryonic stem (ES) cell-derived hepatocytes with introduced *DKC1* mutations revealed aberrant activation of p53 in response to telomere shortening, leading to suppression of HNF4α expression and impaired hepatocyte differentiation ^7^. Counterintuitively, telomere dysfunction and the subsequent p53 response in these ES cell-derived hepatocytes does not elicit apoptosis or senescence, but rather results in hyperproliferation. These results indicate that telomere capping plays an important cell-autonomous role in hepatocytes. However, liver inflammation and cirrhosis are driven in large part via the pathological activation of non-parenchymal cells, specifically the hepatic stellate cells which undergo proliferative expansion and contribute to inflammation and fibrotic scar formation ^21^. Whether telomere dysfunction cell-autonomously affects hepatic stellate cells, or whether telomere dysfunction in hepatocytes promotes a pathogenic response in stellate cells remains unknown.

Here, we model liver phenotypes associated with telomere dysfunction using DC patient-derived iPS cells and isogenic controls with CRISPR/Cas9-mediated homology-directed repair correction of the disease-causing *DKC1* mutation. Differentiation of these cells into hepatocyte-like cells or hepatic stellate cells indicates that the parenchymal hepatocytes are primarily affected by telomere dysfunction. We develop an admixed hepatostellate organoid culture model which further reveals that mutant hepatocytes exert dominant effects on hepatic stellate cells regardless of stellate cell genotype. Hepatostellate organoids containing *DKC1*-mutant hepatocytes exhibit hyperplasia in both the hepatocyte compartment and, remarkably, in stellate cells. Moreover, mutant hepatocytes can induce hallmarks of stellate cell activation independently of stellate cell genotype. Interestingly, mutant hepatostellate organoids also contain off-target PLVAP^+^ endothelial cells reminiscent of scar-associated endothelium observed in non-DC cirrhosis patients ^22^. Ultimately, we demonstrate that inhibition of AKT - a central signaling hub involved in hepatocyte maturation and function - can rescue the liver pathologies in both 2D iPS-derived hepatocytes and 3D hepatostellate organoids. Our findings provide insight into the mechanisms underlying, and potential treatments for liver disease driven by telomere dysfunction.

## Results

### Generation and characterization of DC patient-derived iPS cell-based hepatocyte and hepatic stellate cell models

We initially corrected a *DKC1* A353V mutation (which accounts for approximately 40% of X-linked DC) in DC patient-derived iPS cells using CRISPR-Cas9-driven homology-directed repair (Figure S1A) ^11^. After genome editing, we verified that the resulting isogenic mutant (Mut) and corrected (Cor) iPS cells maintained pluripotency (Figure S1B). To determine whether correction of the mutation restored telomerase function, we measured telomerase activity and telomere length. Corrected iPS cells had higher telomerase activity and longer telomere length compared to isogenic mutants (Figure S1C and S1D).

To model hepatic phenotypes associated with dyskeratosis congenita, iPS cells were differentiated into hepatocyte-like cells (HEPs) using established protocols ^23,24^ with some modification (Figure 1A). At the final stage of differentiation, corrected HEPs exhibited longer telomeres relative to isogenic mutants (Figure 1B). Corrected HEPs exhibited typical hepatocyte morphology (large and polygonal) while mutant HEPs were smaller and had unclear boundaries (Figure 1C). We next assessed hepatic differentiation and function in these cultures. Mutant HEPs had fewer HNF4α-positive (the master hepatocyte transcriptional regulator) and Albumin-positive (ALB) cells, and decreased hepatic function (albumin secretion and low-density lipoprotein uptake) (Figure 1D, S1E, and S1F). As prior studies reported that telomerase mutations can induce steatosis ^14,18^, we also confirmed that mutant HEPs exhibited significantly higher lipid accumulation (Figure S1G). Interestingly, cell-dense, nodule-like structures developed in mutant cultures during hepatic differentiation (Figure S1E and S1H). Given that nodular hyperplasia is reported in human DC patients ^13,25^, we tested whether there was a difference in cell proliferation and found that mutant HEPs had increased proliferation relative to corrected controls (Figure 1E, S1H, and S1I). Taken together, these results indicate that telomerase dysfunction results in disrupted hepatic development and abnormal hepatocyte proliferation. Our findings are largely consistent with a recent study modeling DC phenotypes in human ES-derived hepatocytes ^7^.

**Figure 1.**
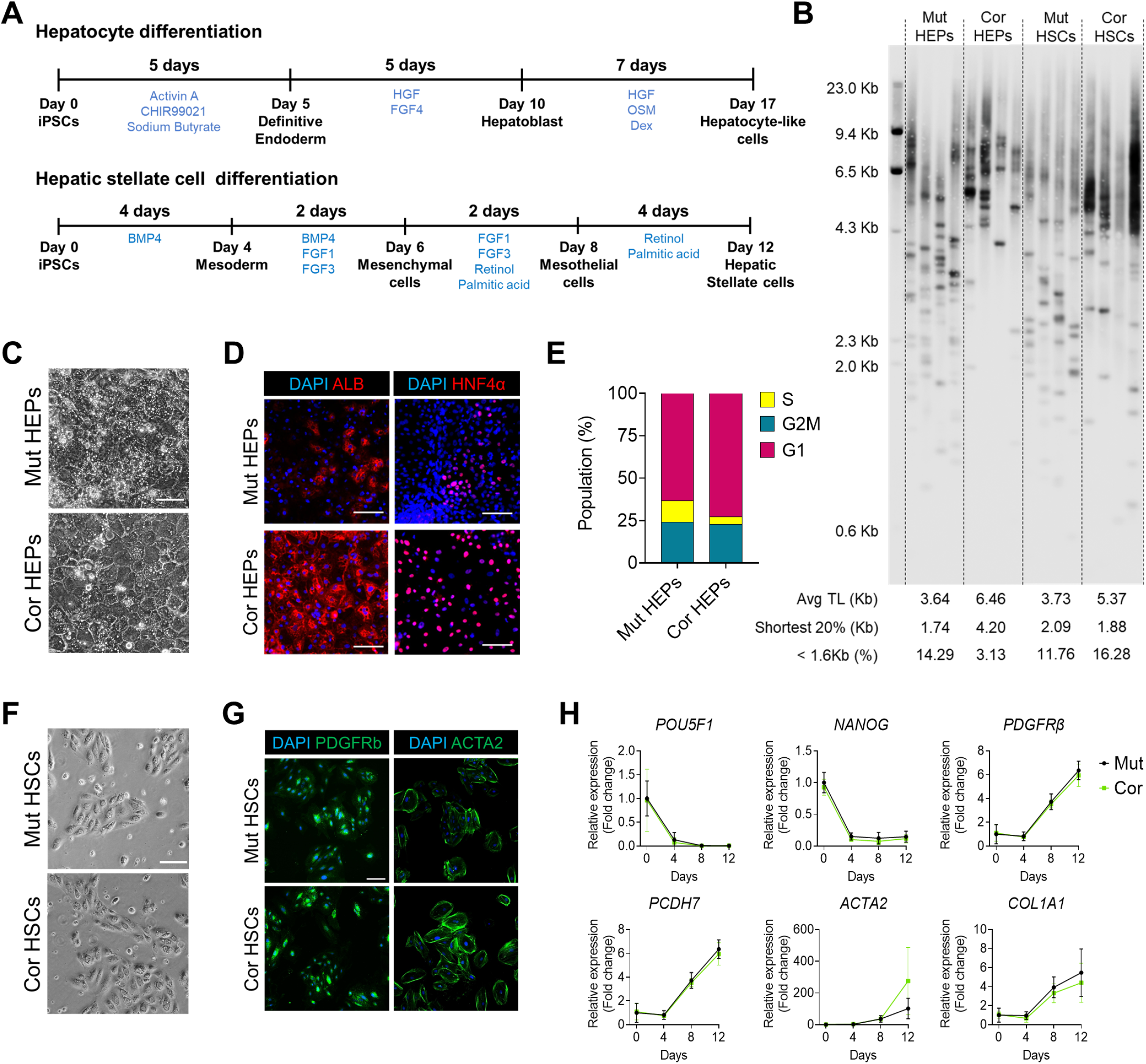
Directed differentiation of HEPs and HSCs from isogenic dyskeratosis congenita iPS cells. (A) Schematic of directed differentiation scheme for iPS cell-derived HEPs and HSCs (B) Telomere length measurement in HEPs and HSCs by Telomere Shortest Length Assay (TeSLA). (C) Representative morphological differences in HEPs on Day 17 (Scale bar = 100 μm). (D) Immunofluorescence images of ALB and HNF4α in HEPs on Day 17(Scale bar = 100 μm). (E) Cell phase analysis of HEPs by Edu assay. (F) Representative morphology of HSCs during the expansion stage (p1) (Scale bar = 100 μm). (G) Staining HSC cultures for stellate cell markers PDGFRβ and ACTA2 (Scale bar = 100 μm). (H) qRT-PCR analysis of pluripotency markers and stellate cell marker genes in HSC cultures. Error bars indicate mean ± SD. See also Figure S1.

We next sought to determine if the *DKC1* mutation also affects the development of non-parenchymal liver cells. We induced differentiation of the isogenic iPS cells into hepatic stellate cells (HSCs) following a recently published protocol (Figure 1A) ^26^. Mutant HSCs had shorter telomeres than corrected HSCs, but these differences were less dramatic than those observed in HEPs (Figure 1B). In contrast to the clear pathologies observed during hepatic differentiation, HSC differentiation revealed no apparent phenotypic differences between the isogenic mutant and corrected cultures (Figure 1F-H). These results suggest that the *DKC1* mutation and consequent telomere shortening primarily effect the parenchymal hepatocytes.

We next performed bulk RNA-seq with hepatoblast (HB) and HEP-stage cultures. Principal-component analysis (PCA) revealed that the primary (PC1) transcriptional differences are driven by differentiation state, while secondary (PC2) differences are driven by genotype (Figure 2A). Gene set enrichment analysis (GSEA) revealed that in both the mutant HBs and HEPs, genes involved in cell proliferation and mRNA translation are highly upregulated (Figure 2B and S2A; Tables S1 and S2). In particular, MYC target genes were highly expressed in mutant cultures, and HNF4α target genes were strongly suppressed in both mutant HBs and HEPs (Figure 2B, S2A, and S2B). Conversely, corrected cultures exhibited enrichment for gene sets related to hepatic differentiation and function. This was confirmed by qRT-PCR, where *HNF4α*, additional hepatocyte nuclear factors, hepatocyte functional markers (*ALB*, *TTR*, and *TDO2*) and apolipoproteins are significantly suppressed in mutant HBs and HEPs relative to isogenic corrected controls (Figure 2C and S2C). We also confirmed that *MYC* and its target genes (including *TERT* and *TP53*) were more highly expressed in mutants (Figure 2C). Immunofluorescent staining revealed an inverse correlation between proliferating cells and differentiated cells in the mutant cultures (HNF4α/FABP1-positive cells are Ki67-negative and vice-versa), suggesting that proliferation and proper hepatocyte differentiation are mutually exclusive in these cultures (Figure 2D, 2E, and S2D). HNF4α is a master regulator of liver development where it regulates target gene expression in a manner antagonistic to MYC ^27^. Our data suggest that the inability of mutant cells to suppress *MYC* expression underlies their failure to activate HNF4α and undergo proper hepatocyte differentiation.

**Figure 2.**
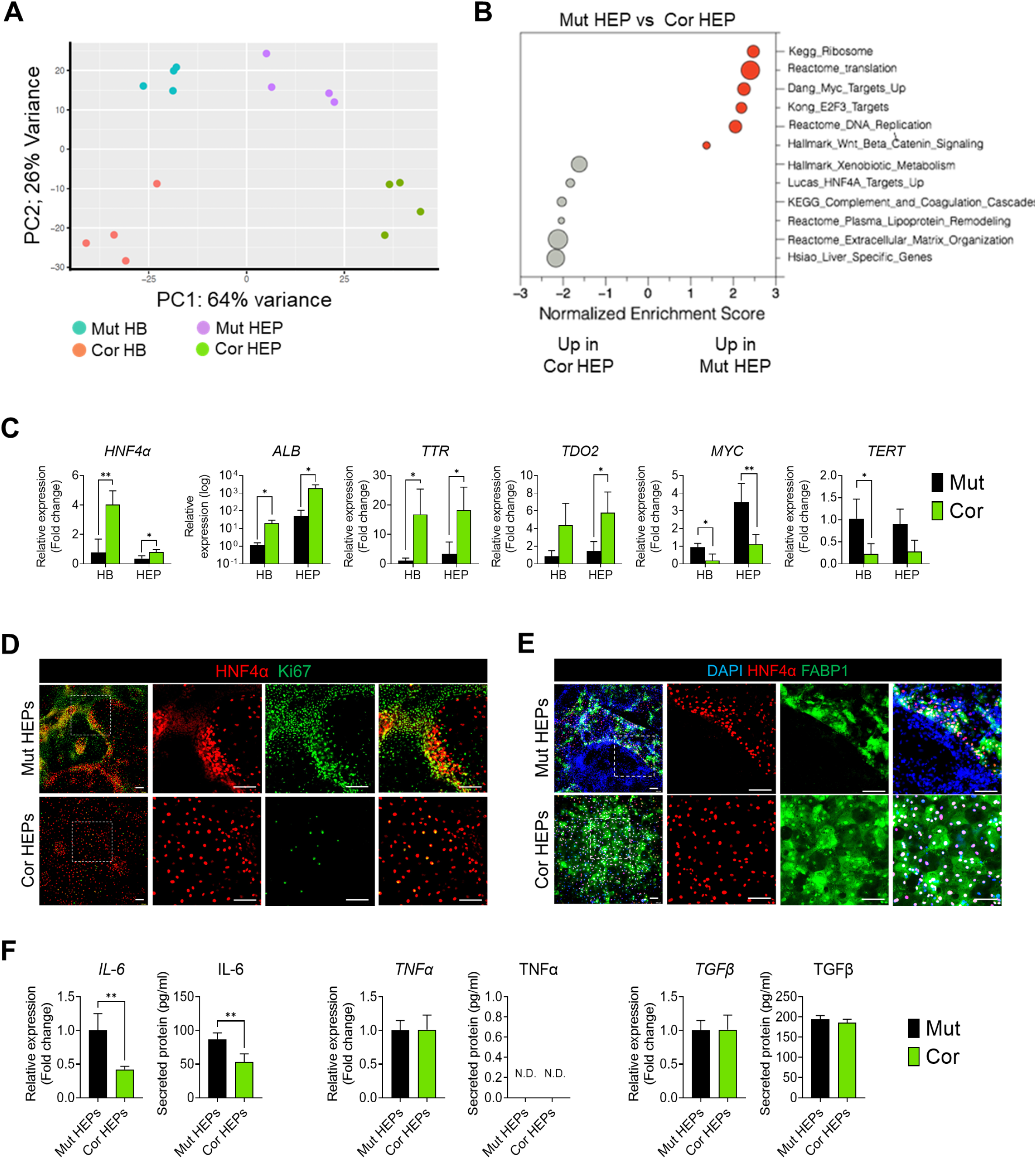
Abnormal hepatic differentiation in *DKC1*-mutant cultures. (A) Comparison of global gene expression in HBs and HEPs by principal component analysis (PCA) of transcriptome profiles (n = 4). (B) Gene set enrichment analysis (GSEA) of differentially expressed genes between mutant HEPs and corrected HEPs. Grey, enriched in mutant HEPs; Red, enriched in corrected HEPs. (C) qRT-PCR analysis of hepatic marker genes (*HNF4α*, *ALB*, *TTR*, *TDO2*), *MYC* and MYC target gene *TERT* (n = 4). *p < 0.05, **p < 0.01. Error bars indicate mean ± SD. (D) HNF4α and Ki67 staining in HEPs (Scale bar = 100 μm). (E) Immunostaining of HNF4α and FABP1 in HEPs (Scale bar = 100 μm). (F) Expression of cytokines in HEPs analyzed by qRT-PCR and ELISA (n = 4). **p < 0.01. Error bars indicate mean ± SD. See also Figure S2, Table S1 and Table S2.

Analysis of pro-inflammatory cytokine expression linked to liver fibrosis revealed high IL-6 expression in mutant HEPs, but no differences in TNFα and TGFβ (Figure 2F). Taken together, these results further confirm that *DKC1* mutation induces abnormal hepatic differentiation and that these abnormal hepatocytes may signal to the microenvironment to promote liver pathologies.

### Modeling DC liver pathologies in hepatostellate organoids

Given that DC mutant hepatocytes exhibit aberrant differentiation and hyperproliferation and produce elevated levels of pro-inflammatory IL-6, we next sought to model the fibrotic and related hepatic phenotypes observed in DC patients that involve non-parenchymal cell types, particularly hepatic stellate cells. We therefore set out to generate ‘hepatostellate’ organoids by admixing HEPs::HSCs or HBs::HSCs at a 5:1 ratio to approximate the hepatocyte::stellate cell ratio found *in vivo* (Fig. 3A-C) ^28^. While HEP::HSC organoids failed to coalesce and grow as 3D structures, HB::HSC mixtures successfully generated 3D hepatostellate organoids followed by final differentiation into mature HEPs. In order to evaluate potential non-cell-autonomous effects of the *DKC1* mutation, we generated organoids composed of HEPs and HSCs of different genotypes (Mut::Cor and Cor::Mut) in addition to mutant-mutant and corrected-corrected hepatostellate organoids (Figure 3A).

**Figure 3.**
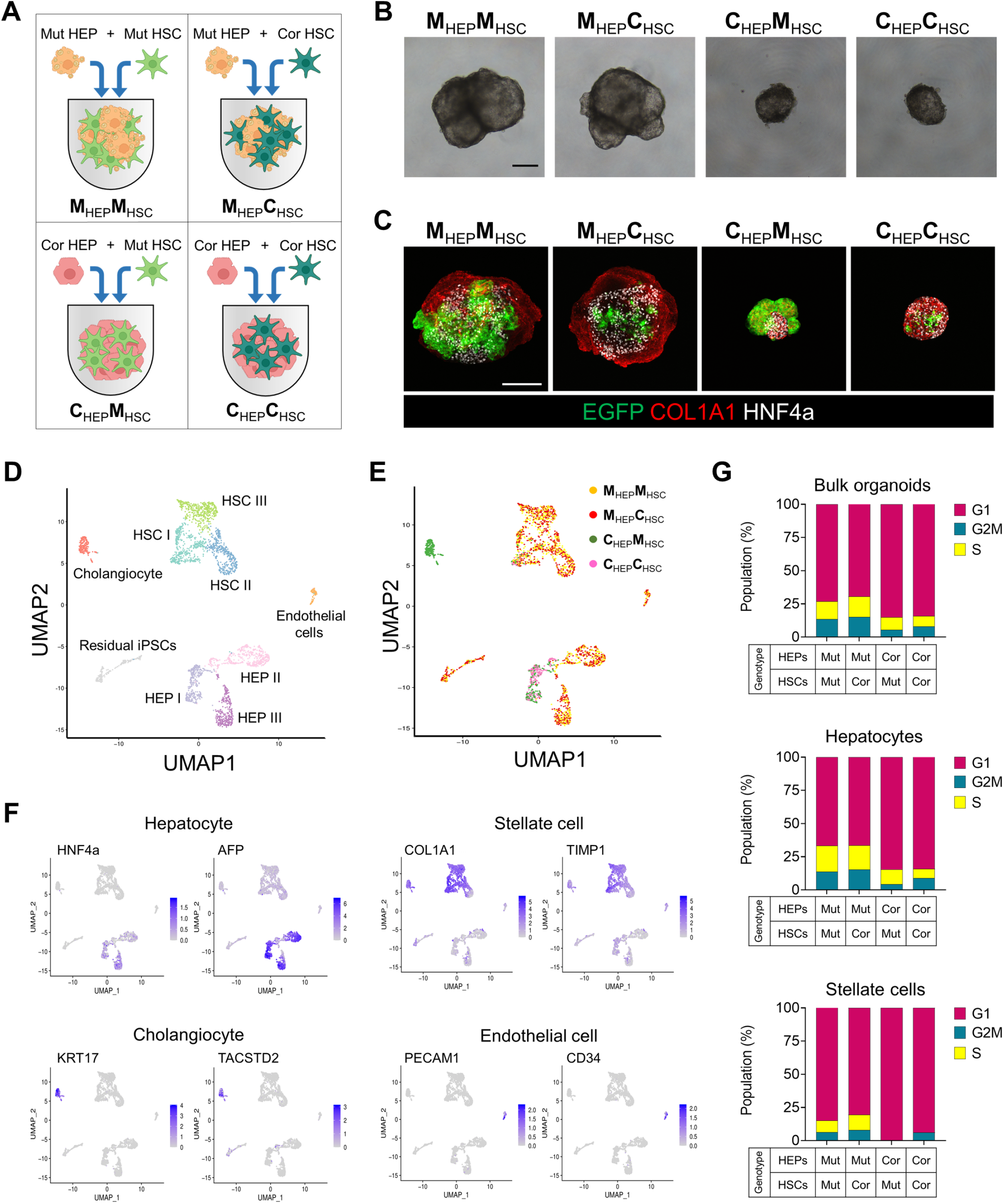
Modeling DC-associated liver phenotypes in hepatostellate organoids. (A) Cartoon depicting the strategy for generating genotype-admixed hepatostellate organoids. (B) Representative morphology of HEP::HSC organoids (Scale bar = 200 μm). (C) Whole-mount immunofluorescent staining of HNF4α and COL1A1 in organoids. EGFP is constitutively expressed from the *AAVS1* locus in the HSCs (Scale bar = 200 μm). (D) Uniform Manifold Approximation Projection or Dimensional Reduction (UMAP) of 2,991 single cell transcriptomes from the four admixed hepatostellate organoid cultures. (E) UMAP annotation by genotypic composition of organoids. (F) UMAPs showing expression of the indicated marker genes for each cell type. (G) Quantification of cell cycle distribution in hepatostellate organoids, and more specifically in hepatocyte and stellate cell clusters, as indicated. See also Figure S3 and S4.

Strikingly, hepatostellate organoid size was dramatically increased in the presence of mutant HEPs, regardless of HSC genotype (Figure 3B, 3C, S3A, S3C, and S3D). To easily distinguish HEPs and HSCs in the organoids, we inserted eGFP into the AAVS1 locus of iPS cells ^29^ and then induced HSC differentiation and hepatostellate organoid formation (Figure S3B). We confirmed that eGFP-expressing HSCs were clearly distinguishable from HEPs, and, interestingly, mutant HSCs appeared consistently larger or more abundant in admixed organoids (Figure 3C). The organoids containing mutant HEPs had more lipid accumulation compared to those containing corrected HEPs (Figure S3E), consistent with phenotypes observed in 2D cultures.

To further characterize the hepatostellate organoids, their cell type-specific gene expression, and the potential for paracrine interactions between HSCs and HEPs, we performed single-cell RNA sequencing (scRNAseq) on the four combinations of admixed organoids (Fig. 3A). Data between replicates was highly congruent (Figure S4A), and these replicates were aggregated for analysis.

Clustering of roughly 3,000 cells from the organoids identified nine populations (Figure 3D). Hepatocytes resided in three clusters (HEP I, HEP II, and HEP III) and expressed classic markers including *HNF4α* and *AFP* (Figure 3D and 3F). Stellate cell markers such as *COL1A1* and *TIMP1* were expressed in three stellate cell clusters (HSC I, HSC II, and HSC III). We also identified a cholangiocyte cluster, an endothelial cell cluster, and some residual undifferentiated pluripotent cells in the organoids (Figure 3D, 3F, and S4B). Cholangiocytes expressed the biliary cell markers *KRT17*, and *TACSTD2* and were found exclusively in corrected HEP::mutant HSC organoids (Figure 3E and 3F). Endothelial cells were identified only in organoids containing mutant HEPs, regardless of HSC genotype, and expressed endothelial cell markers such as *PECAM1* and *CD34* (Figure 3E and 3F). Similarly, residual iPS cells were found almost exclusively in organoids with mutant HEPs, possibly related to their inability to efficiently induce HNF4α and promote proper hepatocyte commitment and differentiation (Figure 3E and S4B). Interestingly, organoids containing mutant HEPs exhibited increased proliferation, not only in the HEP cells, but also in HSCs, regardless of HSC genotype, indicating that *DKC1* mutations in the hepatocytes promote stellate cell activation/proliferation (Figure 3G, S4C, and S4D). Collectively, these findings indicate that the presence of the *DKC1* mutation in HEPs has a dominant effect on the development of admixed hepatostellate organoids.

We next examined the three HEP and HSC clusters more closely (Figure 3D, 4A, and 4C). HEP I is composed primarily of corrected HEPs, regardless of stellate cell genotype (Figure 4A, 4B, and S4E). In contrast, HEP II and HEP III are composed almost entirely of mutant HEPs, and their identity appears unaffected by stellate cell genotype (Figure 4A, 4B, and S4E). HEP I and HEP III express markers of mature hepatocyte differentiation (Figure 4E). In contrast, HEP II is an actively proliferating population with high expression of *MYC* (Figure 4E and S4C). We thus asked what characterizes the difference between the non-cycling ‘wildtype’ HEP I cluster and the ‘mutant’ HEP II/HEP III clusters. Similar to what we observed in 2D HEP cultures, gene set enrichment analysis revealed that cells in the mutant HEP II/III clusters activate MYC targets, along with gene expression programs associated with proliferation and MTORC1 activity (Figure 4G and S4F; Table S3). In contrast, the HEP I cluster where corrected HEPs reside was enriched for gene sets associated with normal liver physiology, such as metabolism of xenobiotics, lipoprotein remodeling, and HNF4α target gene expression (Figure 4G and S4F; Table S3). Thus, *DKC1*-mutant hepatocytes within hepatostellate organoids exhibit hyperplasia and failed terminal differentiation regardless of the genotype of admixed stellate cells.

**Figure 4.**
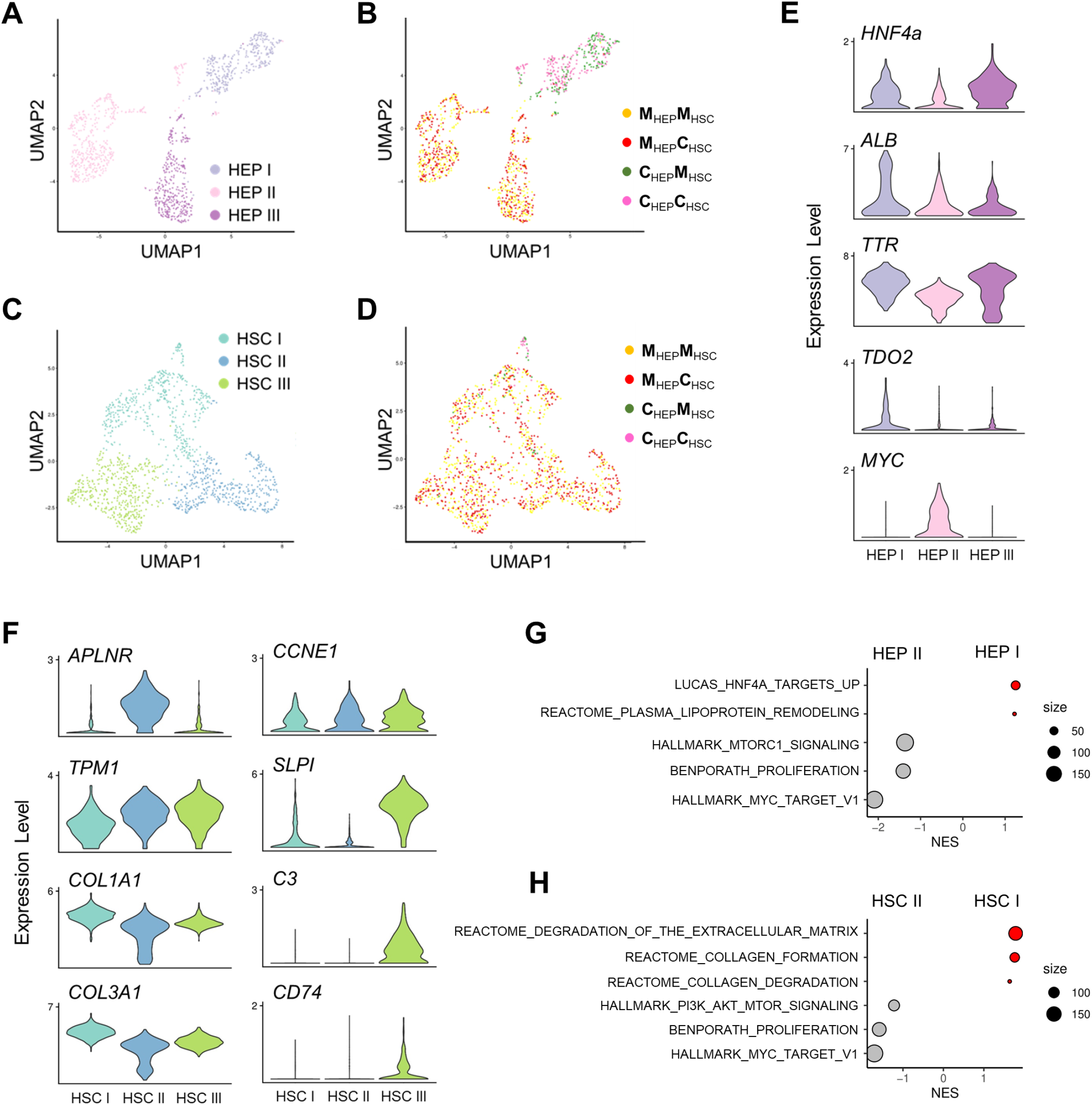
Single cell transcriptomics of hepatostellate organoids reveals *DCK1*-mutant hepatocytes elicit pathological responses in stellate cells. (A and C) HEPs (A) and HSCs (C) from organoids fall into three clusters (I, II, and III) on UMAP projections of single cell transcriptomes. (B and D) Annotation of HEP clusters (B) and HSC clusters (D) by genotypic composition of organoids. (E) Violin plots showing expression of hepatocyte marker genes and *MYC* across three HEP clusters. (F) Expression of activated stellate cell marker genes across three HSC clusters. (G, H) GSEA analysis of HEP I vs. HEP II (G) and HSC I vs. HSC II (H).

In the stellate cell clusters, cells from all admixed organoid conditions could found in HSC I, including all HSCs (both mutant and corrected) co-cultured with corrected HEPs, which are found primarily at the base of the HSC I cluster (Figure 4C, 4D, S4D, and S4E). HSCs cultured with mutant HEPs were rare in our dataset, due to their low input ratio (5 HEPs per HSC) and lack of proliferation (Figure S4D). Cells in clusters HSC II and HSC III were composed exclusively of cells from hepatostellate organoids containing mutant HEPs, further supporting the notion that hepatocyte genotype is the primary driver of HSC phenotype. Relative to HSC II and III, cells in HSC I expressed higher levels genes encoding extracellular matrix components (*COL1A1* and *COL3A1*) (Figure 4F), and were enriched for processes related to the deposition and remodeling of extracellular matrix (Figure S4G). Cells in HSC II (from organoids harboring mutant HEPs) exhibited hallmarks of stellate cell activation. For example, HSC II expressed high levels of *APLNR*, a gene induced in human cirrhotic livers and involved in vascular remodeling ^30,31^ HSC II was also highly proliferative, indicating that the presence of mutant HEPs induces a proliferative response in nearby stellate cells (Figure S4D). Cells in HSC III (also from organoids containing mutant HEPs) also expressed high levels of genes associated with stellate cell activation and inflammation, including *TPM1, SLPI*, *C3*, and *CD74* ^32,33^ Interestingly, both HSC II and HSC III expressed *CCNE1* (Figure 4G), a proliferating cell marker reported to be induced in stellate cells by cMyc-overexpressing hepatocytes in an *Alb-Myc^tg^* mouse model of fibrosis ^34^. GSEA results support this notion, with gene sets associated with PI3K/AKT/MTOR signaling, MYC targets, and proliferation all enriched in HSC II relative to the “wildtype” HSC I cluster where HSCs associated with corrected HEPs reside (Figure 4H; Table S3), independently of HSC genotype. Taken together, these data clearly demonstrate that *DKC1* mutations in hepatocytes play a dominant role in activating stellate cells, by inducing both a proliferative and a pro-inflammatory response associated with hepatic cirrhosis, regardless of stellate cell genotype.

Beyond parenchymal hepatocytes and stromal stellate cells, a recent single cell survey of cirrhotic human livers identified endothelial cell populations associated with fibrotic scarring. These scar-associated endothelial populations were marked by *PLVAP-* or *ACKR1*-positivity and NOTCH ligand expression ^22^. Therefore, we performed additional analysis of organoids containing mutant HEPs where endothelial cells were observed (Figure 5A). Strikingly, these endothelial cells expressed high levels of *PLVAP* and NOTCH ligands *JAG1*, *JAG2*, and *DLL4*, reminiscent of the scar-associated state observed *in vivo* ^22^ (Figure 5B). Hepatostellate organoids expressed high levels of receptors *NOTCH1* and *NOTCH2*, indicating that endothelial-derived NOTCH ligands might act upon HEPs and HSCs in hepatostellate organoids containing mutant HEPs (Figure 5C).

**Figure 5.**
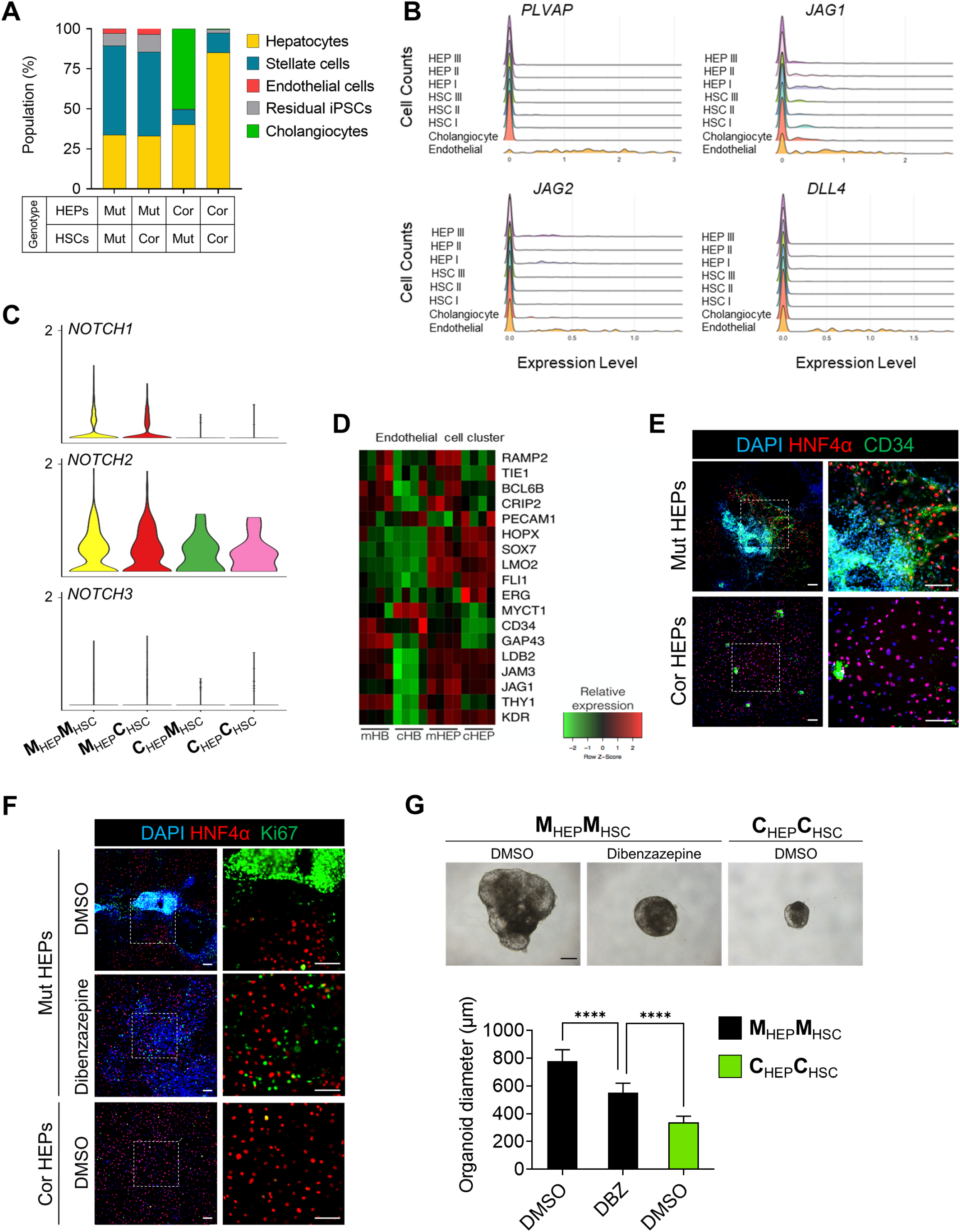
Endothelial cells associated with *DKC1*-mutant hepatocytes exhibit pro-inflammatory phenotypes. (A) Cell composition distribution within hepatostellate organoids. (B) Expression histograms of *PLVAP* and NOTCH ligands (*JAG1, JAG2,* and *DLL4*) in indicated cell cluster from hepatostellate organoid cultures. (C) Violin plot showing NOTCH receptor expression across hepatostellate organoid genotypes. (D) Gene expression heat map visualizing highly expressed genes from the endothelial cell cluster in bulk 2D HB and HEP cultures. (E) Staining of HNF4α and CD34 in 2D HEP cultures (Scale bar = 100 μm). (F) Immunofluorescence images of HNF4α and Ki67 in HEP cultures of indicated genotypes treated with the gamma-secretase inhibitor dibenzazapine (Scale bar = 100 μm). (G) Representative morphology of hepatostellate organoids (Scale bar = 100 μm) and quantification of organoid diameter (n = 20). ****p < 0.0001. Error bars indicate mean ± SD.

Endothelium is traditionally believed to be a mesoderm-derived tissue. In our datasets, endothelial appearance is correlated with mutant hepatocyte genotype, regardless of mesodermal-derived stellate cell genotype (Figure 5A), suggesting that either mutant endodermal hepatocytes instruct endothelial formation within the mesodermal stellate coculture, or that endothelium appears as an off-target cell type in the endodermal hepatocyte directed differentiation. Interestingly, several prior studies reported that endothelial cells can arise during hepatocyte specification from the endoderm, both in human pluripotent-based cultures and in mouse models ^35,36^. Endothelium also plays an important instructive role during liver development ^37^.

We therefore examined our bulk transcriptome datasets from mutant and corrected 2D hepatocyte cultures for evidence of off-target endothelial differentiation and indeed found genes identified in the endothelial scRNAseq cluster to be expressed at higher levels in mutant cultures at both HB and HEP stages versus isogenic controls (Figure 5D). This finding was further supported by staining for endothelial marker CD34 (Figure 5E). Thus, it is most likely that the *DKC1* mutant endodermal hepatocyte cultures promote off-target differentiation of endothelial cells expressing cirrhotic scar-associated genes including NOTCH ligands. Given that activation of the NOTCH pathway has been implicated in liver fibrosis ^38^, we asked whether NOTCH inhibition via the γ-secretase inhibitor dibenzazepine (DBZ) could rescue phenotypes in *DKC1*-mutant hepatocytes and hepatostellate organoids. We found DBZ was able to reduce abnormal nodule formation and organoid size, and reduce but not fully abrogate abnormal proliferation in mutant hepatocytes (Figure 5F and 5G). Taken together these data indicate that endothelial cells may contribute to the liver pathologies in DC patients via NOTCH pathway stimulation.

### Pharmacological rescue of DC-associated live phenotypes

With an aim to more effectively rescue abnormal hepatic differentiation and hepatostellate phenotypes associated with *DKC1-*mutant hepatocytes, we selected several small molecules targeting dysregulated pathways identified in our earlier transcriptomic analyses (Figure 2B, S2A, 4G, 4H, S4F, and S4; Tables S2 and S3). These include inhibitors of GSK3β (a kinase that negatively regulates WNT and MTORC1 pathways) ^39^, and of MYC, AKT, and MTORC1. Of these, the AKT inhibitor MK2206 efficiently inhibited nodule formation, proliferation, and lipid accumulation, along with restoring hepatocyte gene expression (including HNF4α) dose-dependently in mutant HEPs during the differentiation process (Figure 6A-D and S5A). The MYC inhibitor 10058-F4 also inhibited nodule formation, proliferation, and lipid accumulation in mutant HEPs (Figure 6A and 6B). However, it did not re-establish hepatic gene expression as effectively as AKT inhibition (Figure S5B). AKT can negatively regulate GSK3β and positively regulate mTORC1. Interestingly, mTORC1 inhibition with rapamycin was insufficient to suppress MYC expression, and while GSK3β inhibition in combination with AKT inhibition maintained MYC suppression, the addition of GSK3β inhibitor CHIR99021 abrogated the robust HNF4α activation achieved with AKT inhibition alone (Figure S5D) ^40^. Thus, it appears pleiotropic effects downstream of AKT inhibition contribute to the phenotypic rescue observed with MK2206 (Figure S5E).

**Figure 6.**
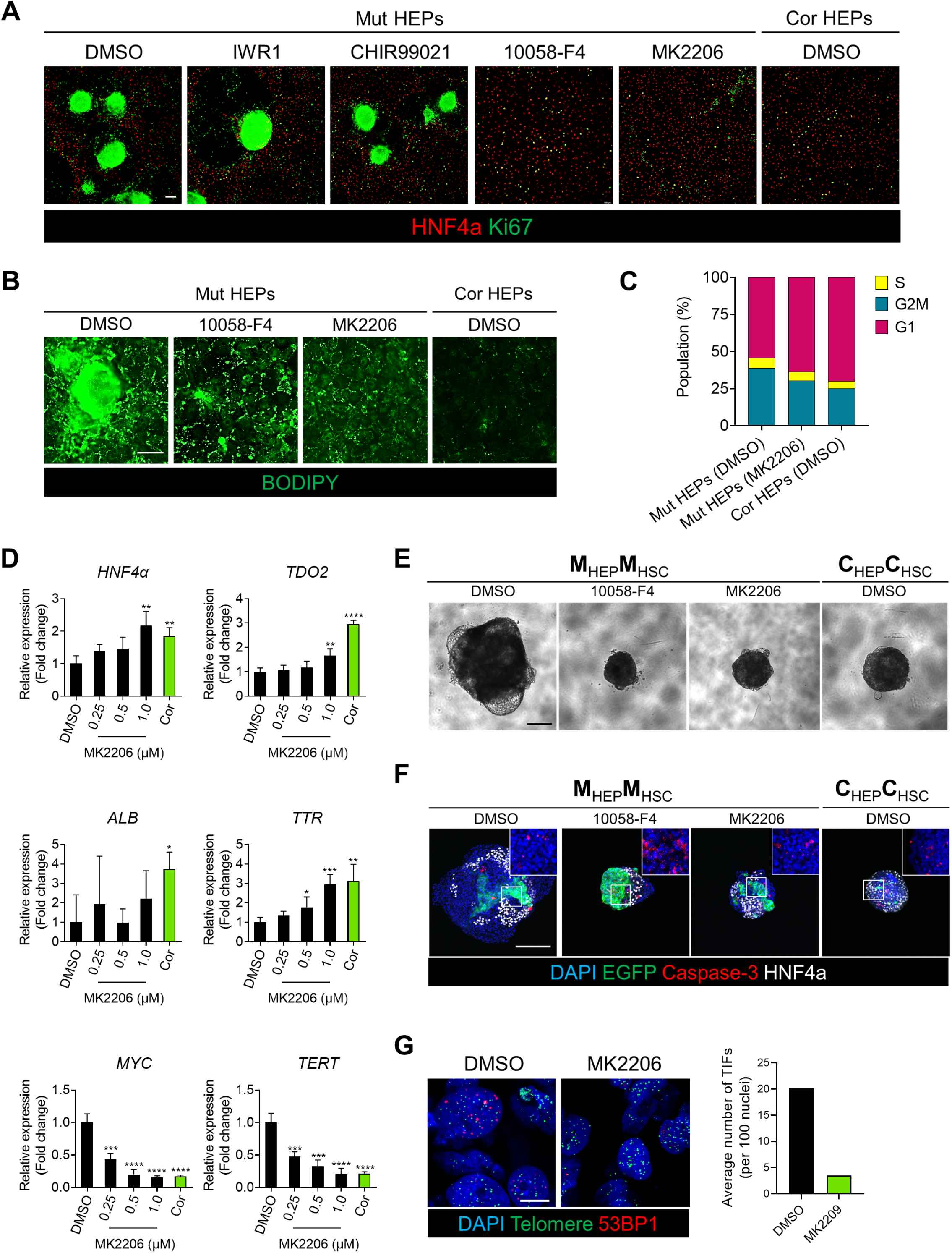
AKT inhibition rescues DC phenotypes in hepatocytes and hepatostellate organoids. (A) Immunofluorescence images of HNF4α and Ki67 in HEPs treated with indicated small molecules (Scale bar = 100 μm). (B) Lipid accumulation in HEPs analyzed by BODIPY staining (Scale bar = 100 μm). (C) Quantification of cell cycle distribution in HEP cultures of indicated genotype and treatment. (D) Expression of hepatic marker genes (*HNF4α*, *TDO2*, *ALB*, and *TTR*), *MYC*, and *TERT* in drug-treated 2D HEP cultures (n = 4). *p < 0.05, **p < 0.01, ***p < 0.001, ****p < 0.0001. Error bars indicate mean ± SD. (E) Hepatostellate organoid morphological changes in response to treatment with MK2206 or 10058-F4 (Scale bar = 200 μm). (F) Whole-mount staining of Caspase-3 and HNF4α in hepatostellate organoids (Scale bar = 200 μm). (G) Representative images showing Telomere Dysfunction Induced Foci (TIF) (Scale bar = 10 μm), quantified at right. See also Figure S5 and S6.

Finally, we generated mutant HEP::mutant HSC organoids using hepatoblasts pre-treated with AKT or MYC inhibitors (MK2206 or 10058-F8) and then induced further hepatic specification (Figure S6A). Organoid size, lipid accumulation, and HNF4α-negative cell populations all decreased after treatment with either drug (Figure 6E, 6F, and S6B-D). However, MYC inhibition with 10058-F8 appeared to increase apoptosis relative to AKT inhibition or control hepatostellate organoids (CorHEP::CorHSC) (Figure 6F). Ultimately, we asked whether the phenotypes observed in mutant hepatostellate organoids and their rescue upon AKT inhibition were correlated with the presence of telomere dysfunction-induced foci (TIFs, defined by colocalization of 53BP1 foci with telomeres). Indeed, we observed TIFs in MutHEP::MutHSC organoids that were markedly reduced upon treatment with MK2206, indicating that AKT inhibition and associated suppression of MYC and activation of HNF4α may be able to feed back to the telomere and promote capping in spite of the *DKC1* mutation (Figure 6G and S6E).

Ultimately, these results indicate that AKT inhibition may represent a viable intervention for preventing DC-associated liver pathologies and demonstrate the utility of the hepatostellate organoid system for modeling these pathologies.

## Discussion

The impact of telomere dysfunction on liver development and physiology in humans is poorly understood. While it is well established that patients with liver cirrhosis or chronic inflammatory conditions exhibit shorter telomeres than healthy age-matched controls, it is unclear how much of this telomere shortening is a consequence of disease, or how much telomere dysfunction may causally contribute to these phenotypes. Recent data suggest a causal role for telomere dysfunction in several liver pathologies: 120 patients with known or suspected telomere disorders (telomeropathies) ^41^ presented with liver involvement in 40% of the cohort, many presenting with hepatomegaly and increased echogenicity ^1^. The fragility of these patients and risks associated with invasive sampling procedures, however, represent significant barriers to tissue acquisition for detailed histological or molecular analyses. In the few instances where biopsied tissue has been analyzed, nodular regenerative hyperplasia, steatohepatitis, steatosis, and cirrhosis are reported ^1,14,42^. These complex liver phenotypes may involve several cell types, including hepatocytes, hepatic stellate cells, and endothelial cells. However, it is less clear which of these distinct cell populations are intrinsically affected by telomere dysfunction, and which are extrinsically responding to signals derived from neighboring cells with telomere dysfunction.

Data in mice support hepatocyte-specific telomere dysfunction as causal in several liver diseases ^20^. An interesting exception is that hepatocyte-specific deletion of *Trf2/Terf2 in vivo* is compatible with normal liver function, which may be explained by the fact that hepatocytes can tolerate the genome endoreduplication that follows the end-end chromosome fusions that re-protect chromosome termini after loss of Trf2. Regardless, this model may not reflect the consequences of telomere uncapping caused by telomere shortening, as happens in DC (Denchi et al., 2006). Mice and humans also exhibit differences in their telomere biology, complicating interpretation of cross-species comparisons. Mice generally express higher levels of telomerase and respond less robustly to telomere dysfunction. Additionally, laboratory mice harbor much longer telomeres than humans necessitating several generations of breeding before exhibiting phenotypes related to telomere dysfunction upon genetic ablation of *Terc* or *Tert* ^43–45^. In contrast, human embryos with complete loss of telomerase activity are not viable, and thus human telomeropathies are generally considered to result from hypomorphic loss-of-function alleles.

Here we utilize isogenic human iPS cells harboring a hypomorphic loss-of-function mutation in *DKC1* which results in telomere dysfunction and dyskeratosis congenita, and their gene-edited normal isogenic controls. The directed differentiation of these lines into hepatocyte-like cells and hepatic stellate cells provides evidence for cell-autonomous phenotypes in hepatocytes, including lipid accumulation and hyperproliferative nodule formation that may be a correlate to the nodular hyperplasia observed *in vivo*^13,25^. Although dyskerin plays roles in ribosome biogenesis, the *DKC1* mutations that cause DC (including the A353V mutation studied here) do not appear to significantly impact this aspect of dyskerin function ^11,12^. Indeed, rather than observing evidence for ribosome-related defects, we instead observed a broad upregulation of genes encoding translation factors in mutant hepatocyte-like cells, consistent with their hyperproliferative nature. Little to no phenotypic changes were observed in stellate cells alone. However, hepatostellate organoids generated by admixing hepatocytes and stellate cells from either *DKC1* mutant or isogenic control cultures revealed a clear instructive role for mutant hepatocytes in inducing a hyperplastic, pro-inflammatory response in stellate cells. Interestingly, off-target endothelial cells expressing genes linked to scar-associated endothelium in cirrhotic human livers ^22^ were observed in cultures containing mutant hepatocytes. Thus, our findings indicate that hepatocytes, and possibly endothelial cells, contribute to the liver phenotypes associated with dyskeratosis congenita, and possibly with telomeropathies more broadly. They further suggest that *DKC1* mutation may also influence aberrant developmental establishment of these tissues, even if overt phenotypes do not manifest until sometime after birth. The notion that telomere dysfunction in hepatocytes may be causal in cirrhotic phenotypes may even extend to idiopathic liver fibrosis, where telomerase mutations are often observed ^16,17^.

Our model ultimately provides a framework for identifying and testing potential interventions in telomere-associated liver disorders. Indeed, inhibition of Notch, which is stimulated by scar-associated endothelial cells and linked to hepatic inflammation and fibrosis ^22,38^ was able to rescue hyperproliferative phenotypes in hepatostellate organoids. Similarly, aberrant AKT activity (a positive regulator of MYC) in hepatocyte-like cells, appears to underly several of the phenotypes we observe both in 2D and hepatostellate organoid cultures. The ability of small molecule inhibitors of AKT to inhibit MYC expression, hyperproliferation and lipid accumulation and restore HNF4α activity may point to a preventative approach to DC-associated liver phenotypes. However, the extent to which such interventions might reverse liver pathologies once they are established remains unclear.

Taken together, organoid-based models of telomeropathies provide a valuable resource for understanding the cellular and molecular basis underlying pathologies in these deadly and largely untreatable diseases. We hope that the hepatostellate organoid model described here provides a framework for vetting future therapeutic approaches targeting the liver pathologies associated with telomere dysfunction.

## Methods

### Cell lines

Male DC patient derived *DKC1* A353V iPS cells were kindly provided by Dr. Timothy S. Olson (Children’s Hospital of Philadelphia). Every iPS cell line (*DKC1* A353V iPS cells, *DKC1* A353V mutation corrected iPS cells, *DKC1* A353V (AAVS1-EGFP) iPS cells and *DKC1* A353V mutation corrected (AAVS1-EGFP) iPS cells) were maintained on vitronectin (Stem Cell Tech.) coated plates using StemMACS™ iPS-Brew XF medium (Miltenyi Biotec) and passaged with 0.5mM EDTA (Invitrogen). iPS cells were cultured at 37 °C in 4% CO_2_, and 5% O_2_ and the medium was replaced every day.

### Correction of *DKC1* mutation in iPS cells

The plasmid, pSpCas9(BB)-2A-GFP (pX458), was obtained from Addgene. Guide RNAs (gRNAs) for correction of *DKC1* A353V mutations were designed using the gRNA design tool from Benchling (https://www.benchling.com/crispr). Designed gRNAs and single stranded DNA donor oligos were synthesized by Integrated DNA Technologies (IDT). gRNAs are inserted into the plasmid and sequenced using the U6 primer (5’-ACTATCATATGCTTACCGTAAC −3’). Patients-derived iPS cells were electroporated with plasmids and single stranded DNA donor oligos using Human Stem Cell NucleofectorTM Kit 1 (Lonza). After 36 hours, transfected cells were selected by GFP positive cell sorting. Sorted cells were plated onto Matrigel (Corning) coated 100mm dishes in StemMACS™ iPS-Brew XF medium with 10μM Y-27632 (Selleck Chemicals). When cell colonies were large enough to be picked, colonies were subcloned and replated into individual wells of Matrigel-coated 96-well plates in StemMACS™ iPS-Brew XF medium with 10μM Y-27632. During replating some iPS cells from each subclone were collected for genomic DNA isolation. After PCR amplification of the targeted region from purified genomic DNA, Sagner sequencing analysis was performed to select *DKC1* mutation-corrected clones.

### HEP differentiation

iPS cells were differentiated into HEPs following previous protocols ^23,24^ with some modification as follows: iPS cells were replated onto Matrigel-coated plates at 0.8×10^5^ cells/cm^2^ in StemMACS™ iPS-Brew XF medium (Miltenyi Biotec) with 10μM Y-27632 (Selleck Chemicals). After 1 day, medium was changed to RPMI-1640 (Lonza) containing 50 ng/ml Activin A (PeproTech), 0.5 mg/ml bovine serum albumin (Sigma-Aldrich), 1x B27 supplement (Gibco), and 0.5 mM sodium butylate (SB, Sigma). The concentration of SB in the medium was the changed to 0.1 mM for 4 days culture in order to induce definitive endoderm differentiation. For the HB induction, definitive endoderm cells were cultured in RPMI-1640 (Lonza) supplemented with 10 ng/mL HGF (Peprotech), and 10 ng/mL FGF4 (Peprotech), 0.5 mg/ml bovine serum albumin (Sigma-Aldrich), and 1x B27 supplement (Gibco) for 5 days. Then, HB stage cells were treated Hepatocyte Culture Medium (without EGF supplement, Lonza) with 10 ng/mL HGF, 10 ng/mL OSM (PeproTech), and 0.1 mM Dexamethasone (Dex). For the inhibition of signaling pathways in differentiating *DKC1* mutant cells, inhibitors were added to HB induction medium (Day 5 – Day 10). Cell cultures were maintained in a 37 °C, 4% CO_2_, and 5% O_2_ environment, and medium was changed every day.

### HSC differentiation and expansion

HSCs were differentiated and maintained as previously described ^26^. iPS cells were seeded onto Matrigel-coated plates and cultured in StemMACS™ iPS-Brew XF medium supplemented with 10 μM Y-27632. Briefly, to prepare HSC differentiation medium, 57% DMEM low glucose (Thermo Fisher Scientific) and 40% MCDB-201 medium (Sigma) were mixed and supplemented with 0.25x linoleic acid-bovine serum albumin (Sigma), 0.25x insulin-transferrin-selenium (Sigma), 1% penicillin streptomycin (Lonza), 100 μM L-ascorbic acid (Sigma), 2.5 mM Dex and 50 mM 2-mercaptoethanol (Thermo Fisher Scientific). On day 0-4, cells were cultured in HSC differentiation medium containing 20 ng/mL BMP4 (R&D) for mesoderm induction. At day 4, growth factors in the medium were changed to 20 ng/mL BMP4, 20 ng/mL FGF1(R&D), and 20 ng/mL FGF3 (R&D). At day 6, differentiating cells were maintained in HSC differentiation medium with 20 ng/mL FGF1, 20 ng/mL FGF3, 5 mM retinol (Sigma) and 100 mM palmitic acid (Sigma). On day 8-12, medium was switched into HSC differentiation medium with 5 mM retinol and 100 mM palmitic acid. Medium was changed every 2 days.

After final differentiation, HSCs were incubated in cell recovery solution (BD) for 30 minutes on ice for recovery. Then cells were harvested using 0.05% Trypsin-EDTA (Life Tech.) and replated on plates coated with Matrigel. Cells were maintained in DMEM high glucose (Thermo Fisher Scientific) with 10% FBS, 5 mM retinol and 100 mM palmitic acid. Medium was refreshed every 2 days. HSCs used in this study were not passaged more than once.

### Immunostaining

Cells were fixed with Paraformaldehyde (PFA) solution 4% in PBS (Santa Cruz biotech.). Fixed cells were permeabilized with 0.1% Triton X-100 in PBS and blocked with 5% serum in PBS. Cells were then incubated overnight at 4 °C with primary antibodies (see antibody list table) and subsequently incubated with secondary antibody (1:500 dilution) for an hour. 4,6-diamidino-2-phenylindole dihydrochloride (DAPI) (Thermo Fisher Scientific) were used for nuclei staining. Stained cells were imaged by LEICA DMI8.

### qTRAP assay

To measure telomerase activity, the qPCR-based telomeric repeat amplification protocol (qTRAP) was performed as previously described ^46^. Briefly, Cells were harvested and resuspended at a concentration of 1,000 cells/μl in NP-40 lysis buffer (1% NP-40). Then, cells were incubated on ice for 30 min and centrifuged for 20 minutes at 4°C. BCA protein assay (Pierce) was performed using the supernatant to determine concentration of the protein. qPCR was conducted using 1μg of lysate. HEK 293 cells were used as telomerase-positive sample for generating a standard curve.

### TeSLA assay

Telomere length was investigated using the telomere shortest length assay (TeSLA) as described ^47^. DNA was extracted using DNeasy Blood & Tissue Kits (Qiagen). DNA concentrations were quantified with a Qubit 2.0 Fluorometer (Invitrogen). 50 ng of each sample were ligated with telorette adaptors. The samples were digested with CviAII, followed by a mixture of BfaI, MseI, and NdeI. The samples were then dephosphorylated and ligated with the TeSLA AT/TA adaptors. 20 ng of each sample were PCR amplified using the FailSafe PCR system with PreMix H (Lucigen). Southern blot-based detection of telomere PCR products was modified from previous protocols ^48,49^. PCR-amplified samples were run on a 0.7% agarose gel at 0.83V/cm for 44 hours at 4°C. The gel was depurinated and denatured, then transferred to a Hybond-XL membrane (Cytiva) overnight with denaturation buffer. The blot was then neutralized with SSC and hybridized with a DIG-labeled telomere probe prepared as described ^48^ in DIG Easy Hyb (Sigma-Aldrich) solution overnight at 42°C. The blot was washed and exposed with CDP-Star (Roche) and visualized in an ImageQuant LAS 4000 (Cytiva). TeSLA quantification was performed using the MATLAB analyzer program, TeSLAQuant.

### TIF assay

Cells were hybridized with a Cy3-labelled PNA telomere repeat probe (Panagene, (5’ -CCCTAA-3’ )) and anti-53BP1 antibodies as described ^50^ with slight modifications. Briefly, 4% paraformaldehyde-fixed cells were permeabilized with PBST (0.2% Triton X-100 in PBS) for 1 min at room temperature, then washed with PBS for 3 min. Cells were then blocked in 4% BSA/PBST for 30 min at room temperature, incubated with rabbit anti-53BP1 antibodies (Novus, NB100-304, 1:100 dilution) for 2 h at 37^0^C in a humidified chamber, washed with PBST, and incubated with Alexa Fluor 647-conjugated goat anti-rabbit antibodies (Invitrogen, A-21244, 1:500 dilution ) for 1 h at 37^0^C. After washing with PBST, cells were incubated with 4% paraformaldehyde in PBS for 20 min to fix antibody in place, followed by quenching of the formaldehyde with 0.25 mM glycine. The cells were subsequently dehydrated with ethanol and air dried. Cy3-conjugated telomere specific PNA probe was applied in hybridization mix (per 100 ul, mix 70 ul of freshly deionized formamide, 15 ul PNA buffer (80 mM Tris-Cl pH 8, 33 mM KCl, 6.7 mM MgCl2, 0.0067% Triton X-100), 10 ul 25 mg/ml acetylated BSA, 5 ul 10 ug/ml PNA probe). Cells were covered with a glass coverslip, denatured at 83 C for 4 min on a heating block and incubated in the dark in a humidified chamber overnight at room temperature. Cells were then washed sequentially with 3 times 70% formamide/2xSSC, 2xSSC and PBS, then blocked and subsequently incubated with Alexa Fluor 647-conjugated donkey anti-goat antibodies (Invitrogen, A-21447, 1:500 dilution), stained with DAPI and mounted in ProLong Gold antifade reagent (Invitrogen, P36935). Confocal images were obtained with the inverted Leica laser-scanning Confocal microscope (TCS SP8), and TIFs were counted by an observer blind to the identity of the samples.

### Quantitative real-time PCR

RNA was isolated using TRIzol™ Reagent (Thermo Fisher Scientific). Total RNA was reverse transcribed using RevertAid First Strand cDNA Synthesis Kit (Thermo Fisher Scientific). qPCR was performed using QuantStudio™ 6 Flex real-time PCR system with power SYBR® green PCR master mix (Applied Biosystems). All gene expression data were normalized by GAPDH using the comparative CT method (2-ΔΔCt). Primer sequences used in this study are listed in Supplementary Table.

### Cell proliferation assay

Ethynyl-2’-deoxyuridine (EdU) incorporation assays were performed to analyze proliferation in HEPs. Briefly, fully differentiated HEPs were cultured with 10μM Edu for 2hours. After 2days, dead cells were labeled with eFluor™ 780 (Thermo Fisher Scientific). Cell proliferation assay were performed using Click-iT Plus Edu Flow cytometry assay kit according to the manufacturer’s instructions. Then fixation, permeabilization and Edu detection were conducted using Click-iT plus Edu flow cytometry assay kits (Thermo Fisher Scientific) according to the manufacturer’s instructions. DNA contents were stained with FxCycle™ Violet (Thermo Fisher Scientific). Cells were analyzed using LSRFortessa flow cytometer (Becton, Dickinson and Company).

### Acetylated-Low-Density Lipoprotein (Ac-LDL) Uptake

After final differentiation, HEPs were cultured with 10 μg/mL 1,1’-dioctadecyl-3,3,3’,3’-tetramethylindo-carbocyaninelabeled Ac-LDL (Thermo Fisher Scientific) at 37 °C for 5 hours. Subsequently, cells were washed with PBS and imaged by LEICA DMI8.

### Western blot analysis

Cells were lysed in RIPA buffer (Abcam) containing protein inhibitor and phosphatase inhibitor cocktail (Cell signaling technology). Protein concentration was estimated by BCA protein assay (Pierce). 30 μg protein was separated by gel electrophoresis, followed by transfer to membranes. Blots blocked with 5% BSA were incubated with primary antibodies for overnight. After washing with TBST, the blots were incubated with HRP-conjugated secondary antibodies. Peroxidase activity was detected by enhanced chemiluminescence substrate (Pierce).

### Accumulated lipid staining

Intracellular lipid accumulation was detected using BODIPY and Oil Red O staining. HEPs and hepatostellate organoids were fixed with 4% PFA solution in PBS (Santa Cruz biotech.). For BODIPY staining, fixed cells or HEPs pre-stained with albumin were incubated with BODIPY 493/503 (1 μg/ml; Thermo Fisher Scientific) and DAPI for 30 min. For Oil Red O staining, fixed HEPs were washed with 60% isopropanol and incubated with 0.5% Oil Red O solution for 30min. To quantify intracellular lipids, extract Oil Red O stain with 100% isopropanol were measured by scanning with microplate reader (OD = 500nm).

### ELISA (ALB and cytokines)

To measure albumin and cytokine secretion level from HEPs, medium was collected after 24 hours (for albumin) or 72 hours (for cytokines) of culture with HEPs and stored at −80 °C until use. Human Albumin ELISA Quantitation Kit (Bethly Laboratory) and IL-6, TNFα, and TGFβ Quantikine ELISA Kits (R&D) were used for ELISA assay according to the manufacturer’s instructions.

### Organoid staining

For whole-mount immunofluorescence staining and imaging of admixed organoids, the organoids were fixed for 30 minutes at room temperature in 4% PFA (Santa Cruz biotech.). After washing with PBS, the organoids were permeabilized in PBS containing 0.5% Triton X-100 for 15 minutes at room temperature, blocked with blocking buffer containing 0.1% Tween 20 and 10% donkey serum (Abcam) for 1 hour at room temperature, and incubated overnight with primary antibodies diluted in blocking buffer at 4°C. Primary antibodies included goat anti-GFP (Abcam), rabbit anti-Ki67 (Abcam), rabbit anti-Collagen I (Abcam), rabbit anti-Cleaved Caspase-3 (Asp175) (Cell Signaling Technology), and mouse anti-HNF-4-alpha (Abcam). The following day, antibodies were removed, and the organoids were washed with PBS containing 0.1% Tween 20. Organoids were then incubated in the dark with secondary antibodies diluted at 1:500 in blocking buffer for 1 hour at room temperature. Nuclei were labeled using DAPI (Thermo Fisher Scientific). For imaging, organoids were mounted on a glass-bottom dish (MatTek) in 1.5% low-melt agarose (Lonza) and cleared overnight using Ce3D ^51^. Fluorescence images were obtained using the inverted Leica laser-scanning Confocal microscope (TCS SP8). Images were processed and brightness and contrast were enhanced using FIJI.

### Generation of EGFP expressing iPS cells

The pX330-SpCas9 and AAVS1-Pur-CAG-EGFP plasmid were acquired from Addgene. gRNAs were designed as previously described ^29^ and cloned into BbsI digested Cas9 plasmid. *DKC1* A353V mutant and corrected iPS cells were transfected with the gRNA-Cas9 vector and the knock-in vector and plated onto plate coated with matrigel. After 24 hours, puromycin selection was performed for additional 3 days to select for positively transfected cells. Surviving colonies were picked and expanded. EGFP knock-in was confirmed by PCR analysis using AAVS1 specific primers ^29^.

### Masson’s trichrome staining

For Masson’s trichrome staining, hepatostellate organoids were fixed in 4% PFA in PBS and transferred into 2% agarose. Then, organoids in agarose gel were embedded in paraffin and sectioned at 4 micron. Collagens in the organoids were stained using Masson’s Trichrome Stain Kit (Polysciences) according to the manufacturer’s protocol.

### Chemical treatment

For rescue assays, Day 5 definitive endoderm stage cells were cultured with inhibitors such as dibenzazepine (10 μM; Selleckchem), IWR-1-endo (10 μM; Selleckchem), CHIR99021 (3 μM; Tocris), 10058-F4 (25 and 50 μM; Cayman), MK2206 (0.25, 0.5, and 1.0 μM; Selleckchem), or dimethylsulfoxide (DMSO) until final differentiation (Day 17). All Inhibitors were dissolved in DMSO.

### mRNA-seq and data analysis

Total RNAs were isolated from HBs (Day 10) and HEPs (Day 17) using TRIzol™ Reagent (Thermo Fisher Scientific) and cleaned up with RNA Clean & Concentrator-5 kit (Zymo Research). Isolated RNAs were submitted to Genewiz sequencing facility for RNA-sequencing. All submitted samples had at least 9.5 of RNA integrity number (RIN). Libraries were prepared by Poly A selection from total RNA and sequenced using HiSeq 2 x 150 bp sequencing (Illumina). FASTQ files from Genewiz were used for further analysis analyses were performed using the statistical computing environment R (v4.0.0), RStudio (v1.1.456), and the Bioconductor suite of packages for R (http://www.bioconductor.org) ^52^. Fastq files were mapped to the human transcriptome (EnsDb.Hsapiens.v86) using Salmon ^53^. Reads were annotated with EnsemblDB and EnsDb.Hsapiens.v86, and Tximport ^54^ was used to summarize transcripts to genes. Limma ^55^ was used to control for batch effects on PCA visualization. For differential gene expression analysis, data were filtered to remove unexpressed and lowly expressed (<10 transcripts across all samples). Differentially expressed genes (false discovery rate < 0.05 and absolute log2 fold change >= 0.59) were identified using DESeq2 ^56^. Heatmaps were created and visualized using gplots. GSEA was used for enrichment analysis ^57^. Raw data is available on the Gene Expression Omnibus (GEO accession number 174018).

### Co-culture organoid induction

HBs at day 10 and fully differentiated HSCs were detached by Accutase (Stem cell tech.) and 0.05% Trypsin-EDTA, respectively. After cell counting using Trypan blue (Thermo Fisher Scientific), HBs and HSCs were mixed in a 5:1 ratio. These mixed cells were seeded into 96 well U bottom plate coated with Nunclon Sphera (Thermo Fisher Scientific) at a density of 3,000 per well. Organoids were cultured in Hepatocyte Culture Medium with 10 ng/mL HGF, 10 ng/mL OSM, and 0.1 mM Dex for 7 days. To inhibit each signaling pathways, inhibitor treated HBs were mixed with HSCs and then inhibitors were additionally treated during the organoid culture period.

### scRNAseq

Hepatostellate organoids at day 17 were treated with dissociation buffer, (1:1:1 mixture of 0.25% Trypsin-EDTA, Accutase (Stem cell Tech.), and Collagenase IV (1 mg/mL, Gibco)) at 37 °C for 20min. Cells were filtered through a 40 micron Flowmi cell strainer (Bel-Art) before flow sorted using Aria B (Becton, Dickinson and Company). Gating was applied using the FACSDiva software (Becton, Dickinson and Company) to remove dead cells and aggregates. Sorted cells were encapsulated using the Chromium Controller (10x Genomics) and the Chromium Single Cell 3’ Library & Gel Bead Kit (10x Genomics) following the standard manufacturer’s protocols. In brief, cells loaded onto the Chromium controller were limited to 10,000-12,000 to reach a multiplet rate no higher than 6%. All libraries were quantified using an Agilent Bioanalyzer and pooled for sequencing on an Illumina NovaSeq at Center for Applied Genomics (CAG) at Children’s Hospital of Philadelphia. At least two technical replicates were run in parallel for each sample. Targeted median read depth is 50,000 reads per cell from total gene expression libraries and 10,000 reads per cell for hashtag barcode libraries. CellRanger (version 3.1.0) was used to align reads to the GRCh38-3.0.0 transcriptome and quantify read and hashtag counts ^58^. Seurat (version 4.0.1) was used for standard QC, hashtag doublet removal, log normalisation, regression of difference between S and G2M phase cells and multiple clustering techniques (PCA, TSNE, UMAP) following the appropriate Seurat workflows and their parameters ^59^. VisCello (version 1.1.1) was used for visualization of cell clusters, differential expression, gene ontology and KEGG pathway analyses using default parameters ^60^. PHATE clustering was performed using the PhateR (version 1.0.7) package ^61^.

## QUANTIFICATION AND STATISTICAL ANALYSIS

All data were expressed as the arithmetic mean ± standard deviation of the mean (SD). Statistical analysis was performed with GraphPad Prism 9 (GraphPad Software) using Student’s t test. P-values less than 0.05 were considered statistically significant. Statistical parameters, including numbers and significance, are shown in the legend for each figure.

## Data availability

The accession number for the RNA-seq data reported in this paper is GEO: GSE 174018. Single Cell RNAseq data are currently being deposited.

## Acknowledgments

We thank members of the Johnson and Lengner labs, as well as Dr. Foteini (Faye) Mourkioti and the Mourkioti lab for discussions on the study and manuscript. We also thank Team Telomere for their continued support of this research. We thank the Cell and Developmental Biology (CDB) Microscopy Core for access to the confocal microscopy equipment used in our work. This work was supported by NIH grants R01HL148821 (F.B.J. and C.J.L), R21AG054209 (F.B.J. and C.J.L.) and a grant from Team Telomere and the Penn Orphan Disease Center. This work was also supported in part by the National Institute of Diabetes and Digestive and Kidney Diseases (NIDDK) Center for Molecular Studies in Digestive and Liver Diseases (P30DK050306) and its core facilities.

## Author contributions

YJC conducted all experiments unless otherwise indicated below. YJC contributed to experimental design, interpretation, and writing of the manuscript. MSK assisted with cell culture and hepatostellate culture and performed confocal imaging and analyses. JHR, NMJ, NL, and YJC contributed to the design, execution, and analysis of single cell transcriptomics. CTB, NL, and YJC contributed to the design, execution, and analysis of bulk transcriptomics. SR performed telomere Southern blotting. QC performed TIF assays. YT performed cell proliferation assays. RJF, SAT, and YJC contributed to generating of the cell lines. ZC performed western blot analysis. FBJ and CJL contributed to study design, experimental analyses, and writing of the manuscript.

## Supplemental Figure Legends

**Supplementary Figure 1. Generation of iPS cell lines and analysis of HEPs after directed differentiation**. (A) Sanger DNA sequencing tracks showing correction of the *DKC1* mutation in patient-derived iPS cells. The PAM sequence for the RNA was mutated by employing a GCT codon at A353 instead of the wild type codon (GCG) for the correction. (B) Immunostaining of pluripotency markers Nanog and Sox2 in the isogenic iPS cell pair (Scale bar = 100 μm). (C) Telomerase activity in iPS cells measured by quantitative telomeric repeat amplification protocol (qTRAP) assay (n = 3), **p < 0.01. Error bars indicate mean ± SD. (D) Telomere length in iPS cells measured by Telomere Shortest Length Assay (TeSLA). (E) Acetylated-low-density lipoprotein (Ac-LDL) uptake assay in HEPs (Scale bar = 100 μm). (F) Albumin secretion by HEPs was analyzed using ELISA (n = 4), **p < 0.01. Error bars indicate mean ± SD. (G) Lipid accumulation in HEPs analyzed by BODIPY staining and Oil Red O staining. Scale bars, 100 μm. ***p < 0.01. Error bars indicate mean ± SD. (H) Ki67 and E-cadherin staining in HEPs. White arrows indicate nodule-like structures (Scale bar = 100 μm). (I) EdU cell proliferation assays using flow cytometry analysis.

**Supplementary Figure 2. Comparison of HBs and HEPs derived from isogenic DC iPS cells**. (A) Gene set enrichment analysis (GSEA) was performed to identify gene sets enriched differently in mutant HBs and corrected HBs. (B) Heatmaps showing differences in gene expression within indicated gene sets during hepatic differentiation. (C) Quantitative RT-PCR analysis of hepatocyte markers and *TP53*. (n = 4). *p < 0.05, **p < 0.01, ***p < 0.001. Error bars indicate mean ± SD. (D) HNF4α and Ki67 staining of mutant HBs and corrected HBs (Scale bar = 100 μm).

**Supplementary Figure 3. Characterization of hepatostellate organoids.** (A) Quantification of hepatostellate organoid diameter (n = 15). ****p < 0.0001. Error bars indicate mean ± SD. (B) Images of EGFP expression after targeting to the endogenous *AAVS1* locus in iPS cells, and in iPS cell-derived HSCs (Scale bar = 100 μm). (C) Masson’s Trichrome Stain in organoid sections (Scale bar = 100 μm). (D) Images of whole-mount staining of COL1A1 and HNF4α (Scale bar = 100 μm). (E) BODIPY staining images of organoids (Scale bar = 200 μm).

**Supplementary Figure 4. Single cell transcriptomics in hepatostellate organoids.** (A) UMAP annotated by experimental replicate. (B) Expression of pluripotency (*NANONG, POU5F1*) and definitive endoderm (*SOX9*, *EPCAM*) marker gene expression. (C and D) Cell cycle phase analysis in hepatocyte clusters (C) and stellate cell clusters (D). (E) The fraction of cells that contributed to each of the three HEP and HSC clusters by organoid genotype. (F, G) GSEA analysis of HEP I vs. HEP III (G) and HSC I vs. HSC III (H). See also Table S3.

**Supplementary Figure 5. Pharmacological rescue of DC phenotypes in HEP cultures.** (A) Flow cytometry plots of EdU cell assays. (B) Gene expression analysis of hepatic markers and *MYC* in 2D HEP cultures treated with MYC inhibitor 10058-F4 (n = 4). *p < 0.05. Error bars indicate mean ± SD. (C) Immunofluorescence staining for HNF4α and Ki67 in 2D HEP cultures treated with indicated small molecules (Scale bar = 100 μm). (D) Western blot analysis of proteins involved in the AKT signaling pathway. (E) Schematic illustration of AKT signaling in hepatocytes.

**Supplementary Figure 6. Pharmacological rescue of DC phenotypes in hepatostellate organoids.** (A) Schematic of drug treatment period during organoid induction. (B) Quantification of organoid diameter in response to indicated drug treatment (n = 20). ****p < 0.0001. Error bars indicate mean ± SD. (C) Whole-mount immunofluorescent staining of HNF4α and Ki67 in organoids. (Scale bar = 200 μm) (D) BODIPY staining of hepatostellate organoids. The scale bar represents 100 μm. (E) Representative images showing 53BP1 and γ-H2A.X staining in HEP cultures (Scale bar = 100 μm).

