## Supplementary Figures for "Patient induced pluripotent stem cell-derived hepatostellate organoids establish a basis for liver pathologies in telomeropathies"

Young-Jun_Supplementary Figure 1

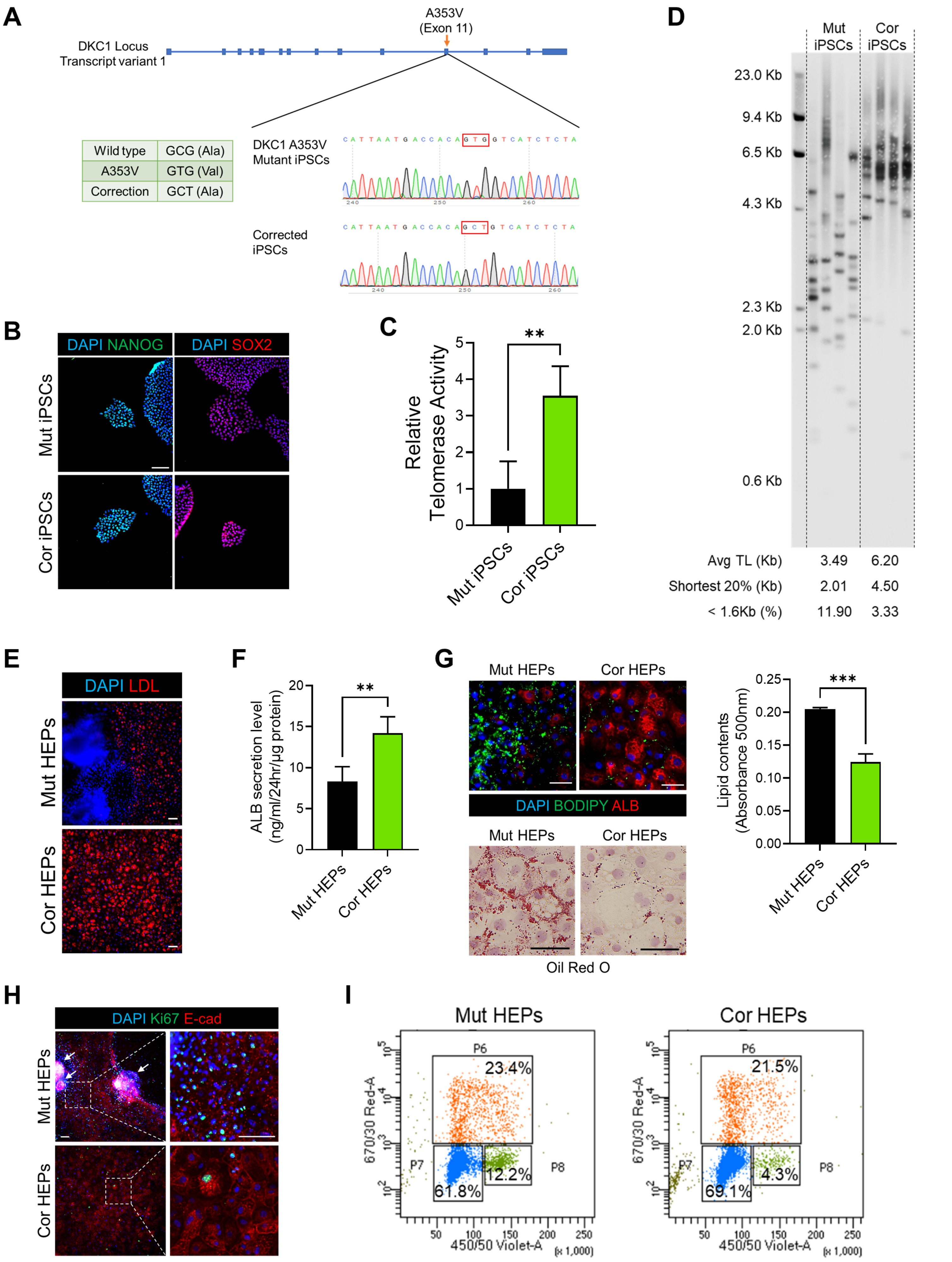

Young-Jun_Supplementary Figure 2

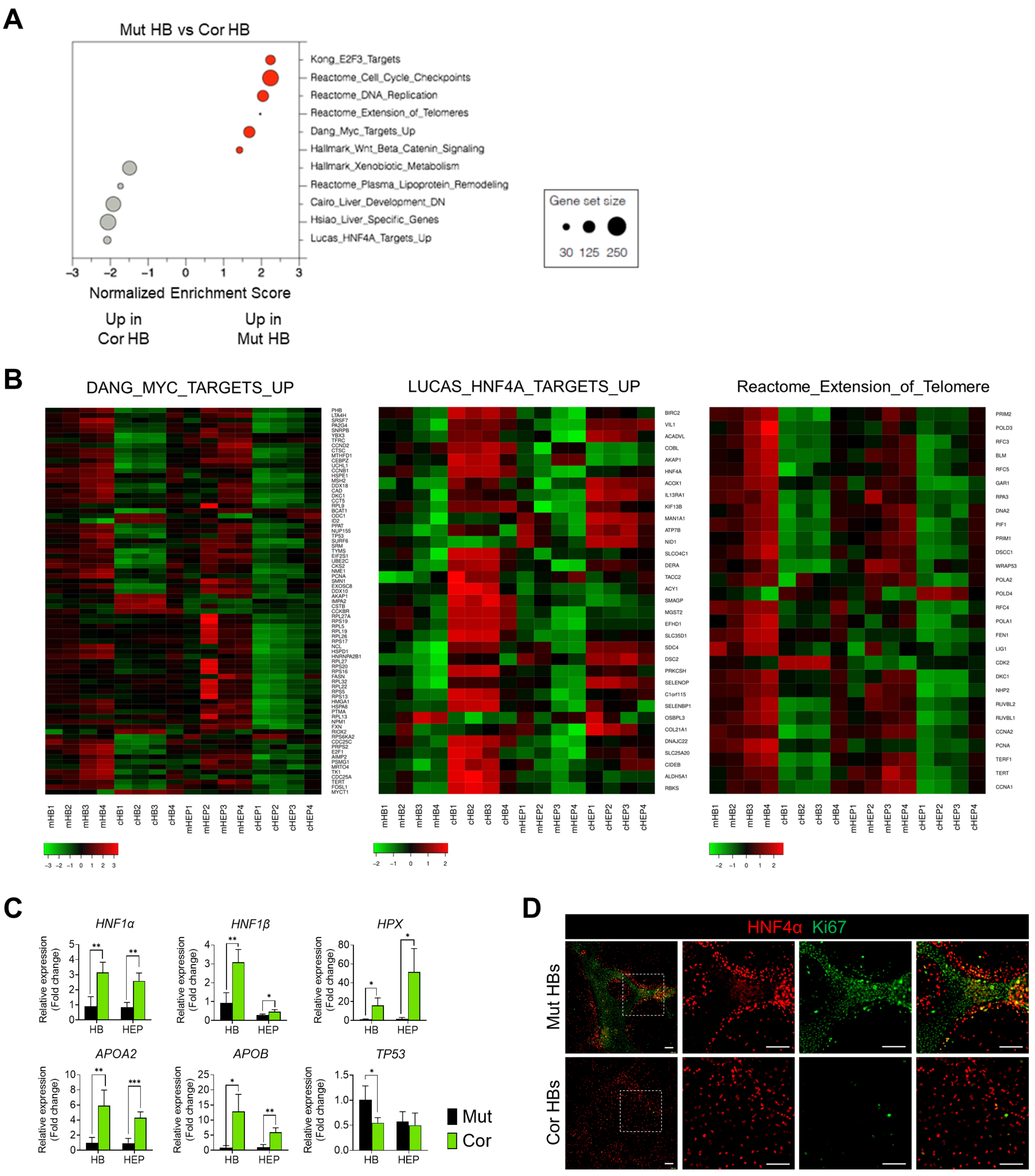

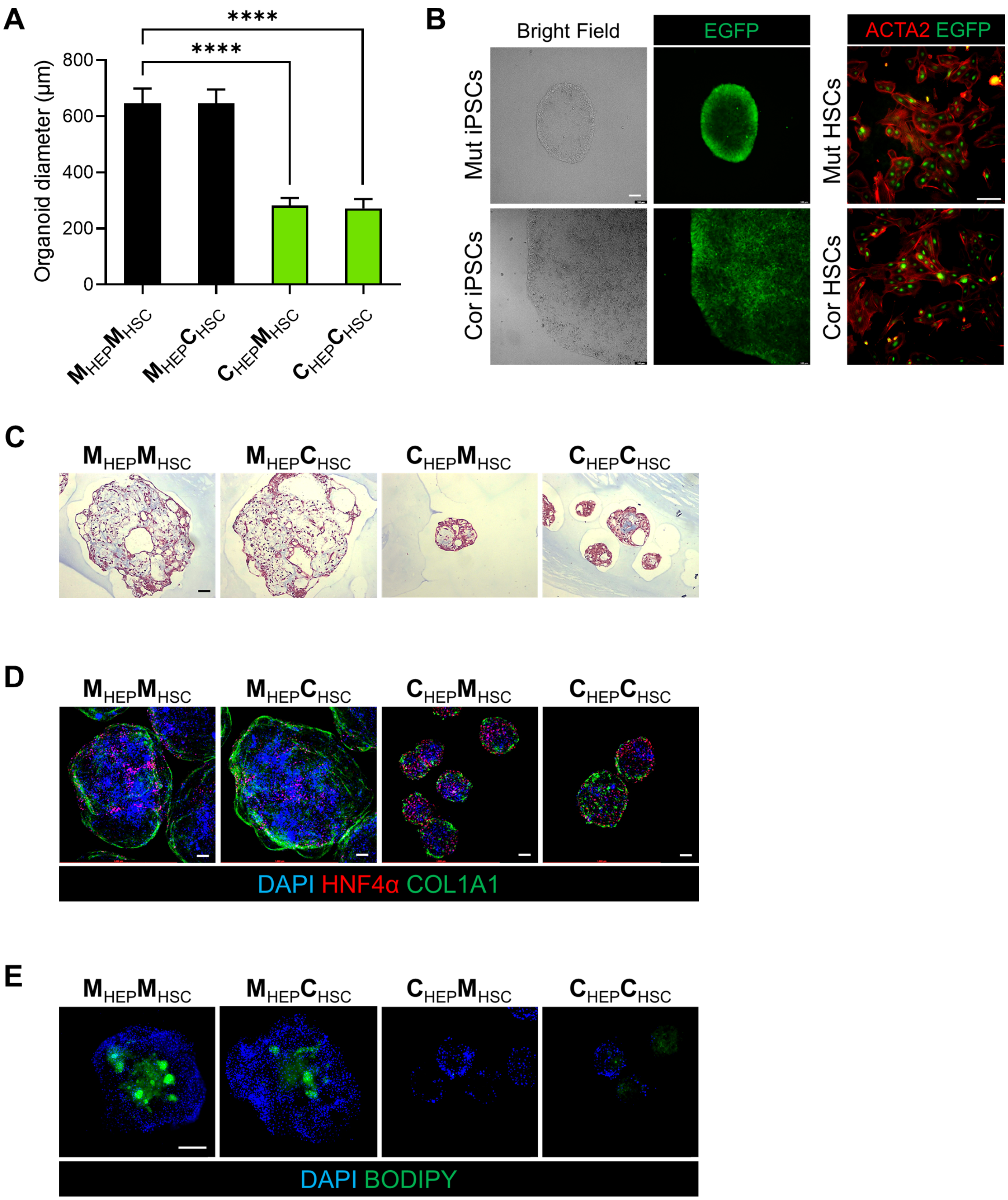
A
B
C
D
E
****
****
Organoid diameter (μm)
800
600
400
200
0
MHEPMHSC
MHEPCHSC
CHEPMHSC
CHEPCHSC
Bright Field
EGFP
ACTA2 EGFP
Mut iPSCs
Cor iPSCs
Mut HSCs
Cor HSCs
DAPI HNF4α COL1A1
DAPI BODIPY

Young-Jun_Supplementary Figure 4

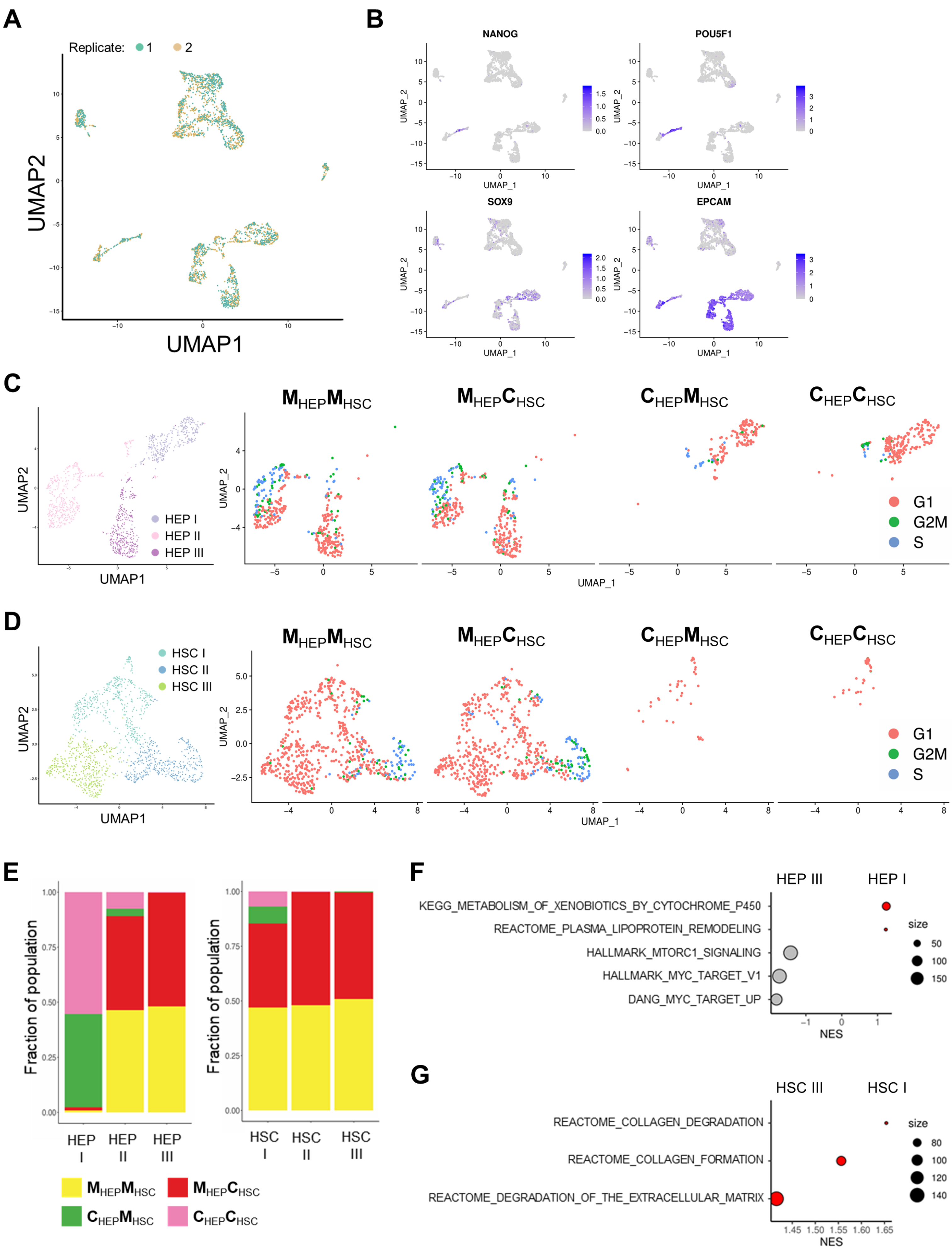

### Young-Jun_Supplementary Figure 5

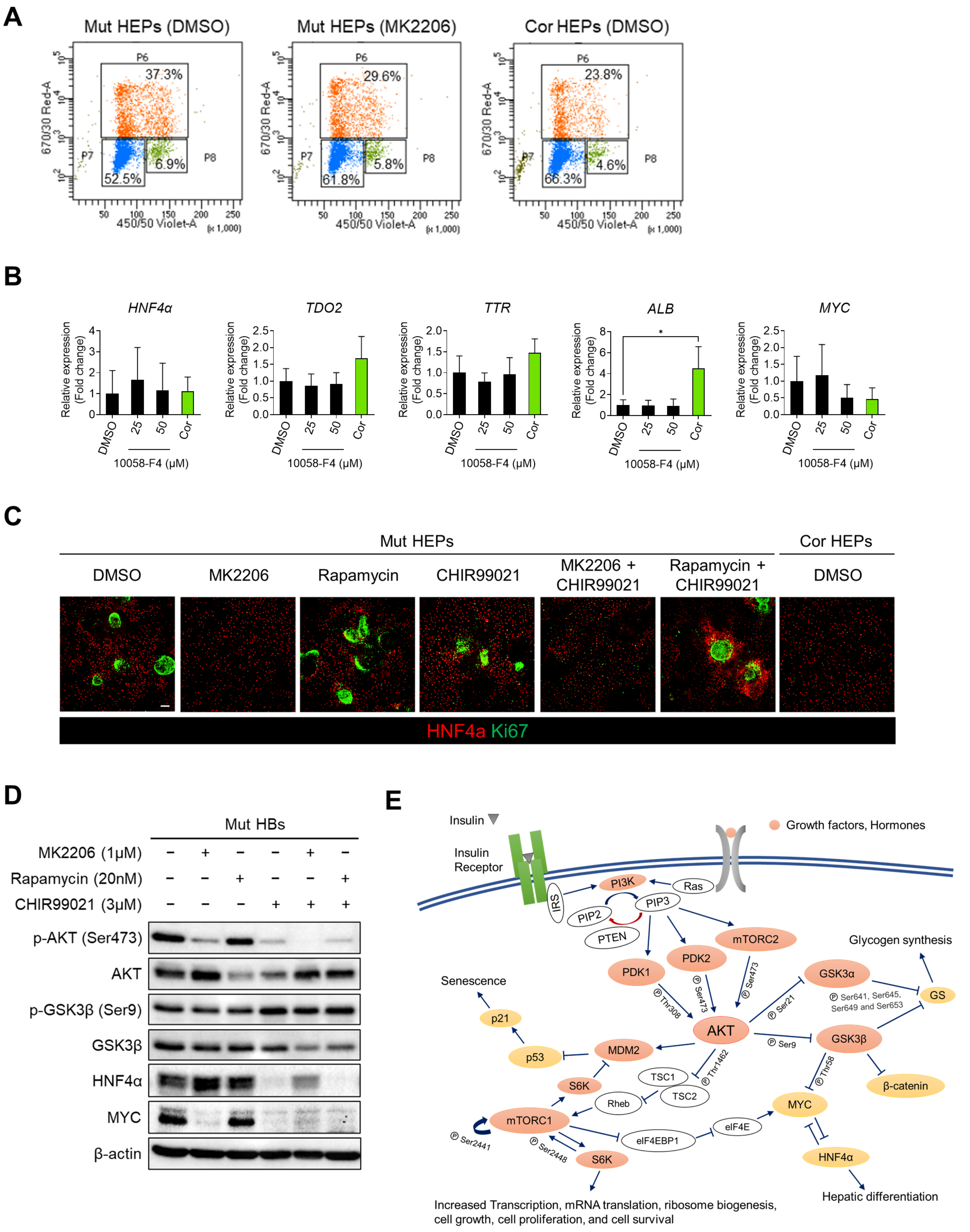

Young-Jun_Supplementary Figure 6

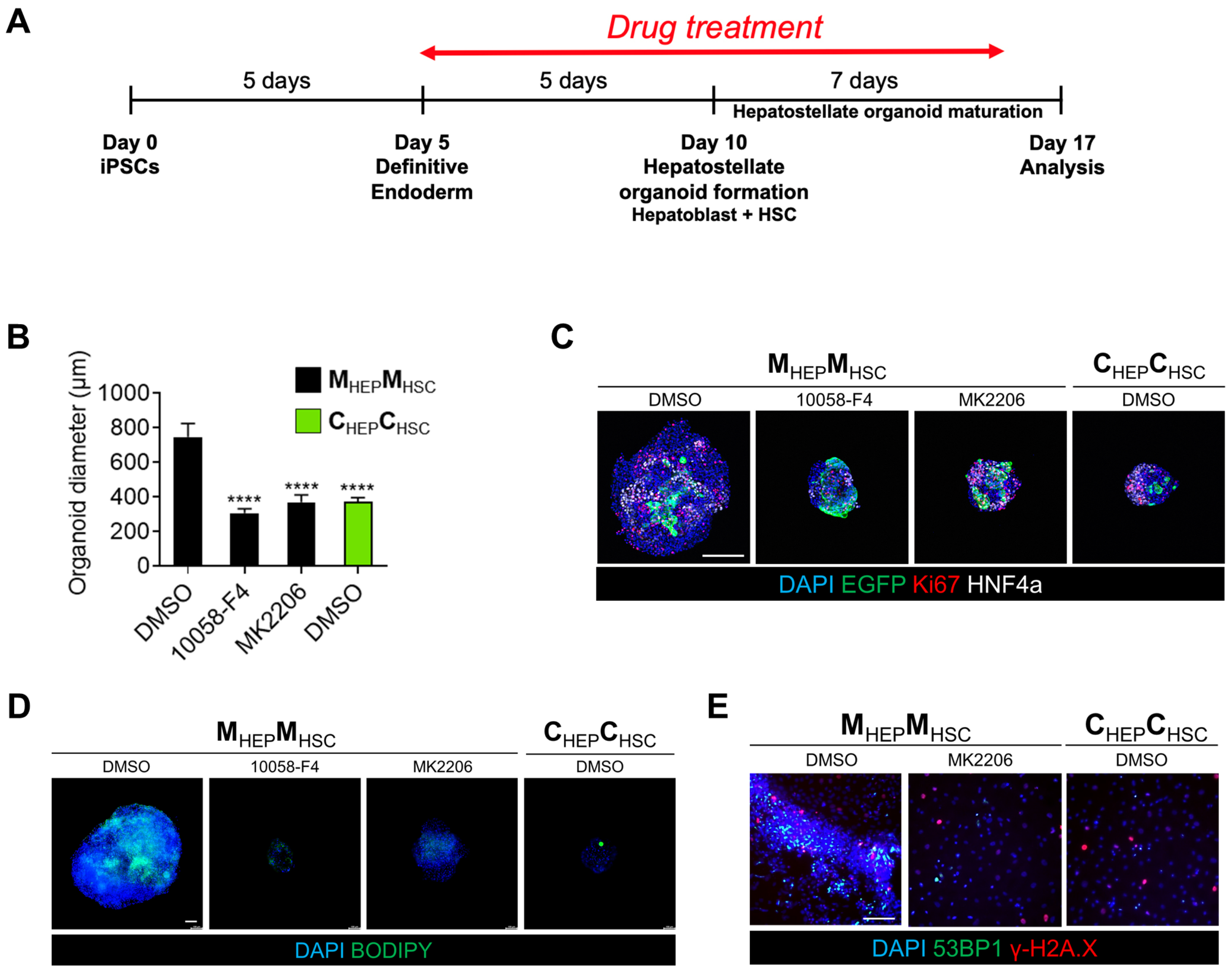
